# Activation of PKR by a short-hairpin RNA

**DOI:** 10.1101/2024.05.08.592371

**Authors:** Kyle A. Cottrell, Sua Ryu, Helen Donelick, Hung Mai, Jackson R. Pierce, Brenda L. Bass, Jason D. Weber

**Affiliations:** Department of Medicine, Division of Molecular Oncology, Washington University School of Medicine, Saint Louis, Missouri, USA; Department of Cell Biology and Physiology, Washington University School of Medicine, Saint Louis, Missouri, USA; Department of Biology, Siteman Cancer Center, Washington University School of Medicine, Saint Louis, Missouri, USA; ICCE Institute, Washington University School of Medicine, Saint Louis, Missouri, USA; Department of Biochemistry, Purdue University, West Lafayette, IN, USA; Department of Biochemistry, University of Utah, Salt Lake City, UT, USA

**Keywords:** double-stranded RNA, dsRNA, PKR, RNA interference, RNAi

## Abstract

Recognition of viral infection often relies on the detection of double-stranded RNA (dsRNA), a process that is conserved in many different organisms. In mammals, proteins such as MDA5, RIG-I, OAS, and PKR detect viral dsRNA, but struggle to differentiate between viral and endogenous dsRNA. This study investigates an shRNA targeting DDX54’s potential to activate PKR, a key player in the immune response to dsRNA. Knockdown of DDX54 by a specific shRNA induced robust PKR activation in human cells, even when DDX54 is overexpressed, suggesting an off-target mechanism. Activation of PKR by the shRNA was enhanced by knockdown of ADAR1, a dsRNA binding protein that suppresses PKR activation, indicating a dsRNA-mediated mechanism. In vitro assays confirmed direct PKR activation by the shRNA. These findings emphasize the need for rigorous controls and alternative methods to validate gene function and minimize unintended immune pathway activation.

## Introduction

Recognition and response to viral infection are essential processes in all kingdoms of life. For organisms infected by viruses, recognition of viral infection is often dependent upon sensing double stranded RNAs (dsRNA) generated from the viral genome. In mammals, multiple sensor proteins detect viral dsRNA. The RIG-I like receptors MDA5 and RIG-I, activate the type I interferon (IFN) pathway following detection of viral dsRNA ^1^. Oligoadenylate synthase (OAS) proteins activate RNase L ^2^, while protein kinase R (PKR) activates the integrated stress response pathway after sensing viral dsRNA ^3^. Each of these pathways relies on a sensor protein binding to dsRNA; however, these proteins cannot distinguish between viral dsRNA (exogenous) and dsRNAs arising from inter- and intramolecular base-pairing between endogenous RNAs ^4^.

Endogenous dsRNAs arise from a variety of RNA transcripts. While some endogenous dsRNAs are formed through base-pairing interactions between repetitive elements, such as inverted Alu repeats ^5^, other, often much shorter, endogenous dsRNAs are intermediates of the microRNA (miRNA) biogenesis pathway ^6^. Primary miRNA (pri-miRNA) transcripts are generally transcribed by RNA polymerase II and form stem-loop structures with mismatches, loops and bulges. These pri-miRNAs are processed by DROSHA and its partner protein DGCR8 in the nucleus to form stem-loop structures of between ∼55-70 nucleotides long, referred to as precursor miRNAs (pre-miRNA). Finally, the pre-miRNA is processed further in the cytoplasm by DICER and its accessory proteins to form mature miRNAs that are loaded on to an argonaute protein to form the miRNA-induced silencing complex (miRISC). The components of miRNA processing, specifically DICER, also processes exogenous short-hairpin RNAs (shRNAs) ^7^. By using the cellular machinery associated with miRNA processing and miRNA-mediated repression, shRNAs are routinely used to knockdown the expression of genes of interest. While shRNAs can provide robust knockdown of target genes, they have been plagued by off-target effects, usually through miRNA like effects on non-target mRNAs ^8-10^.

The protein double-stranded RNA dependent protein kinase (PKR) serves an important role in sensing viral dsRNA. PKR is the primary mechanism of the innate immune system to trigger apoptosis and inhibit protein synthesis ^11-13^. PKR contains three domains: two dsRNA binding domains (dsRBD), dsRBD1 and dsRBD2, and a kinase domain ^14^. Activation of PKR requires dsRNA binding, dimerization, and autophosphorylation ^15^. Upon binding of dsRNA by two PKR monomers, PKR dimerizes, bringing the kinase domains of each monomer close enough to autophosphorylate and activate PKR ^16,17^. Active PKR has several canonical substrates, including the alpha subunit of eukaryotic initiation factor 2 (eIF2α), which when phosphorylated by PKR inhibits translation ^18^. Key to PKR activation are its dsRBDs. Each dsRBD is capable of binding dsRNA in a sequence independent manner, and can bind dsRNAs as short as 16 bp ^19,20^. While PKR has an important role in sensing viral dsRNAs, because the dsRBDs bind in a sequence-independent manner PKR can also bind numerous endogenous dsRNAs ^21^. Here, we provide evidence for a shRNA that directly activates PKR in human cell lines.

## Results

### Knockdown of DDX54 causes activation of PKR

Using proximity labeling, we had previously identified the RNA helicase DDX54 as an ADAR1 interacting protein ^22^. This led us to study the role of DDX54 in the regulation of dsRNA sensing, a role that has been well established for ADAR1. To begin to evaluate the role of DDX54, we utilized lentiviral shRNA mediated knockdown by RNAi to reduce DDX54 expression in two ADAR1-dependent cell lines. These cells require ADAR1 for viability and will activate multiple dsRNA sensing pathways following genetic depletion of ADAR1 ^23^. One specific shRNA sequence (encoded within two shRNA expression plasmids from The RNAi Consortium (TRC) library, TRCN0000049957 and TRCN0000289044, referred to here as shDDX54-4 and shDDX54-5, though the plasmids encoding these shRNAs are slightly different, the shRNA itself is identical in sequence), showed substantial knockdown of DDX54 expression, Figure 1a. Interestingly, we observed robust activation of PKR, as indicated by PKR phosphorylation at Thr446 and phosphorylation of eIF2alpha at Ser51, in cells transduced with lentivirus encoding shDDX54-4 and shDDX54-5, Figure 1a-b. In addition to activation of PKR, we observed reduced cellular proliferation, and/or viability, as indicated by reduced foci formation, Figure 1c-d. These observations were consistent with the effect of ADAR1 knockdown in these same cell lines ^23^.

**Figure 1:**
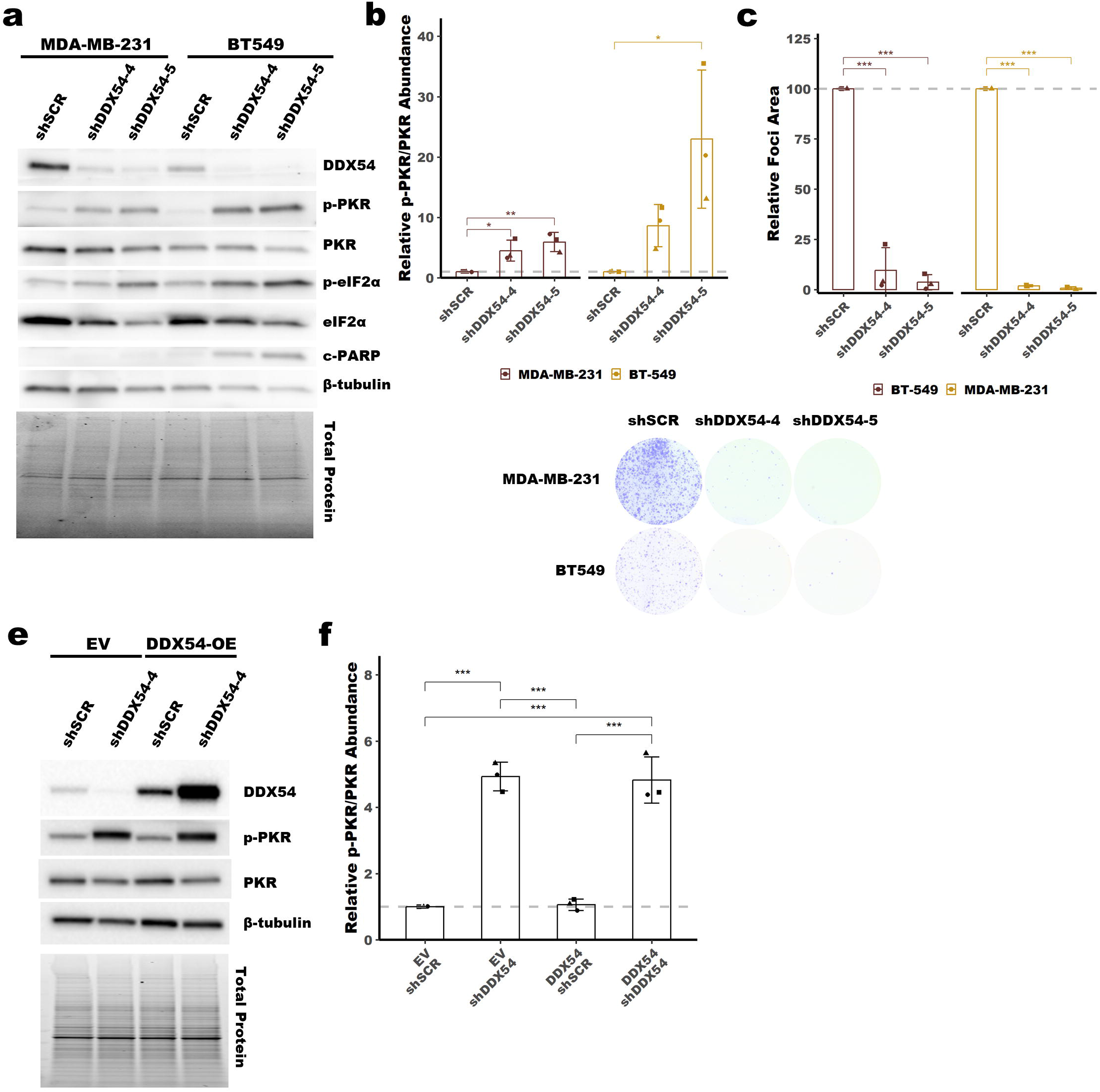
An shRNA targeting DDX54 causes activation of PKR. **a** Representative immunoblot of pPKR and other proteins of interest following knockdown of DDX54 by shDDX54-4 and shDDX54-5 (identical shRNA sequence) in MDA-MB-231 and BT549. Total protein was imaged using a Stain-Free Gel and was used for normalization. **b** Quantification of pPKR/PKR from the immunoblot in **a**. See Supplemental Figure 1 for quantification of other proteins of interest. **c** Quantification of foci formation shown in **d. e** Representative immunoblot of pPKR and other proteins of interest following knockdown of DDX54 by shDDX54-4 in empty vector (EV) control BT549 or DDX54 overexpressing (DDX54-OE) BT549. Bars represent the average of at least three replicates (shown as differently shaped points), error bars are +/- SD. ^*^ p <0.05, ^**^ p <0.01, ^***^ p < 0.001. P-values determined by Dunnett’s test **b-c** or one-way ANOVA with post-hoc Tukey **d**.

### PKR activation is an off-target effect of the shRNA targeting DDX54

Based on the DDX54 knockdown phenotype, we began to explore the mechanism for PKR activation following knockdown of DDX54 in BT549. To verify that the effects we had observed were caused by reduced DDX54 expression, and to rule out off-target effects of the shRNA, we performed a rescue experiment. We overexpressed a wobble mutant of DDX54 (not targetable by shDDX54-4) and knocked down endogenous DDX54 using shDDX54-4. Surprisingly, we observed that exogenous DDX54 expression was not sufficient to prevent activation of PKR in cells expressing shDDX54-4, Figure 1e-f. This finding suggested that shDDX54-4 caused PKR activation through an off-target mechanism, and not through reduced expression of DDX54.

### Knockdown of ADAR1 increases PKR activation in combination with the shRNA targeting DDX54

To narrow down the mechanism of PKR activation in cells expressing shDDX54-4, we combined expression of shDDX54-4 with knockdown of ADAR1 in two ADAR1-independent cell lines. These cell lines were chosen because we have previously observed that neither activate dsRNA sensing pathways following knockdown of ADAR1 or DHX9 alone – two proteins shown to suppress activation of dsRNA sensors ^23,24^. We hypothesized that if the effect of shDDX54-4 on PKR activation was dsRNA mediated, that knockdown of ADAR1 could potentially increase PKR activity because of ADAR1’s known role in suppression of PKR activation through binding to dsRNA. In support of this hypothesis, we observed that ADAR1 knockdown enhanced PKR activation in cells expressing shDDX54-4, Figures 2ab and 2ef. Additionally, knockdown of ADAR1 and expression of shDDX54-4 caused reduced foci formation (Figure 2c-d and 2g-h). The effects of ADAR1 knockdown on PKR activation in cells expressing shDDX54-4 are consistent with increased activation of PKR due to increased abundance of dsRNA in cells expressing shDDX54-4.

**Figure 2:**
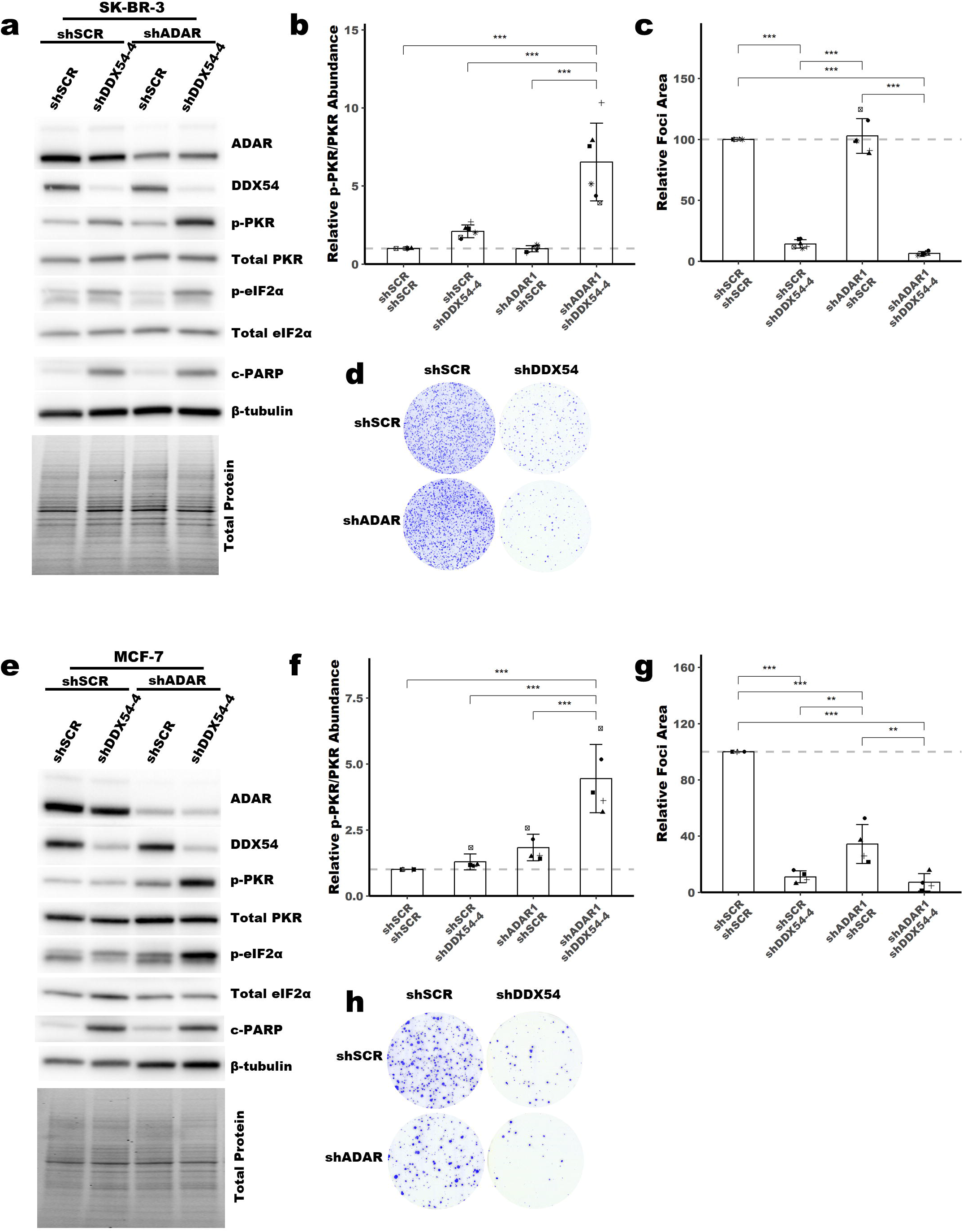
Knockdown of ADAR1 enhances PKR activation in cells expressing shDDX54-4. **a** Representative immunoblot of pPKR and other proteins of interest in SKBR3 expressing shDDX54-4 or shSCR, with or without knockdown of ADAR1. Total protein was imaged using a Stain-Free Gel and was used for normalization. **b** Quantification of pPKR/PKR from the immunoblot in a. See Supplemental Figure 2 for quantification of other proteins of interest. **c** Quantification of foci formation shown in **d** for SK-BR-3. **a** Representative immunoblot of pPKR and other proteins of interest in MCF-7 expressing shDDX54-4 or shSCR, with or without knockdown of ADAR1. Total protein was imaged using a Stain-Free Gel and was used for normalization. **b** Quantification of pPKR/PKR from the immunoblot in **a**. See Supplemental Figure 2 for quantification of other proteins of interest. **c** Quantification of foci formation shown in **d** for MCF-7. Bars represent the average of at least three replicates (shown as differently shaped points), error bars are +/- SD. ^*^ p <0.05, ^**^ p <0.01, ^***^ p < 0.001. P-values determined by one-way ANOVA with post-hoc Tukey.

### shRNA targeting DDX54 activates PKR in vitro

Based on the above data, we hypothesized that shDDX54-4 (and shDDX54-5 which are identical in sequence) may directly activate PKR. To evaluate this, we performed an in vitro kinase assay with purified PKR and shDDX54-4 or shSCR control. As a positive control, we also performed the assay with 106 BLT RNA previously shown to robustly activate PKR ^25^. Unlike shSCR, shDDX54-4 activated PKR in a dose-dependent manner and showed substrate inhibition at higher concentrations as previously observed for PKR, Figure 3a-b ^26^. These findings are consistent with shDDX54-4 directly activating PKR in cells expressing the shRNA.

**Figure 3:**
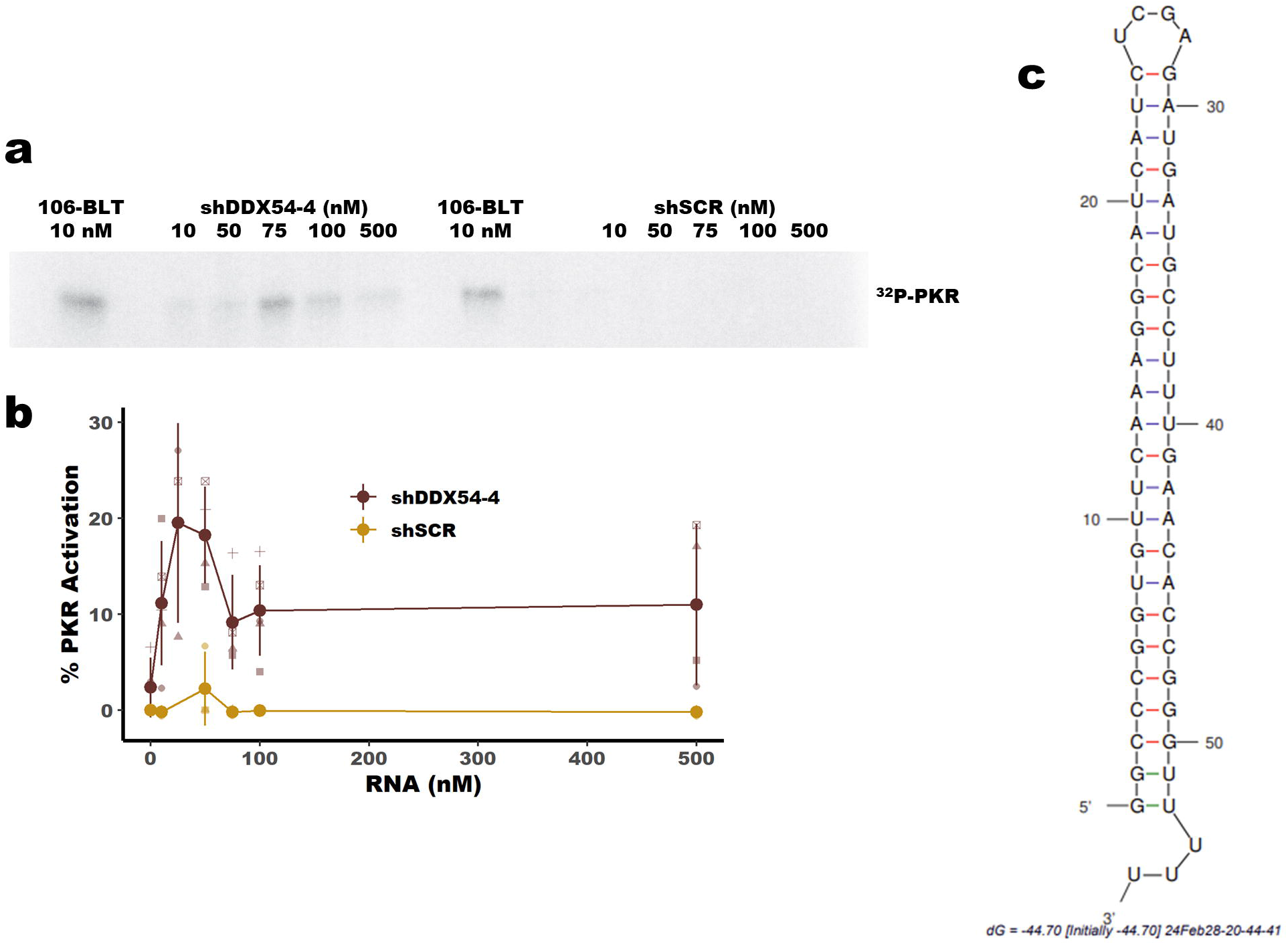
**a** Representative PhosphorImage of PKR phosphorylation following in vitro kinase assay with RNAs indicated. **b** Quantification of PhosphorImage shown in **a**. Larger points represent the average of at least three replicates (shown as differently shaped smaller points), error bars are +/- SD. **c** Predicted structure of shDDX54-4 as determined by mFold ^39^.

## Discussion

Activation of dsRNA sensors, such as PKR, has become a promising therapeutic strategy for cancer. For example, depletion of ADAR1, a suppressor of dsRNA sensing, has been shown to promote cell death and anti-tumor immunity in many cellular and in vivo models of cancer ^23,27-29^. Many of those studies, and others that focus on different suppressors of dsRNA sensing, have utilized shRNA mediated knockdown to reduce the expression of genes of interest ^24,30^. Here, we report the direct activation of PKR by an shRNA targeting DDX54. We observed activation of PKR by this shRNA in cells constitutively expressing the shRNA and in an in vitro PKR kinase assay. Importantly, our in vitro kinase activity showed substate inhibition of PKR kinase activity, consistent with previous in vitro studies of PKR activation ^26^. The finding that an shRNA can directly activate PKR brings to light a challenge for studies focused on dsRNA sensing. It is possible, without the correct controls, that experiments that utilize shRNAs may lead to false-positive data indicating that a given gene plays a role in suppression of dsRNA sensing.

Our finding of a shRNA activating PKR is not entirely unique. Early in the use of siRNAs, many groups observed off-target activation of PKR. Off-target activation of PKR has been observed for several different siRNAs targeting multiple genes ^31-35^. Those findings and our own are complicated by biochemical studies of PKR activation that have revealed the requirements for activation of PKR by dsRNA. In vitro studies have shown that a dsRNA of at least 33 bp is required to recruit two PKR monomers and activate PKR, but each dsRBD can independently bind 16 bp length dsRNA ^20^. Recent studies suggest that the minimal length of dsRNA required to activate PKR may change under certain circumstances. Several natural RNA activators of PKR fall below the 33 bp length requirement for PKR activation, such as influenza B ribonucleoprotein (16 bp dsRNA) and interferon-γ mRNA (5-7 bp dsRNA pseudoknot) ^36,37^. Another short RNA element that has been shown to activate PKR is the human immunodeficiency virus 1 (HIV-1) Tat-responsive region RNA (TAR) ^38^. The TAR RNA forms a hairpin structure of similar size to an shRNA, 23 bp, but normally inhibits PKR activation. However, TAR can form intermolecular dimers of sufficient length to activate PKR. Thus, PKR is activated by shorter dsRNAs through multiple mechanisms.

The question remains as to how the shRNA identified here directly activates PKR, while many other shRNAs used by our laboratories and countless other laboratories do not activate PKR. One intriguing aspect of the shRNA is the GC rich sequence near the base of the stem (Figure 3C). The GC rich region includes seven G-C base pairs, which presumably would stabilize the duplex and thus may contribute to PKR activation. Whether the GC rich stem of the shRNA is important for PKR activation remains to be determined by future experiments.

In the meantime, it is important for researchers to utilize appropriate controls to mitigate the effects of direct activation of PKR (and potentially other dsRNA sensors) by shRNAs. Important measures that can be taken include, 1) Using multiple shRNAs, 2) Performing rescue experiments, 3) Using an orthogonal approach to reduce gene expression, such as knockout by CRISPR-Cas9 or using CRISPR inhibition. It should be noted that when we began these experiments, we did attempt to use multiple shRNAs. We ordered two different shRNA clones from Millipore-Sigma (TRCN0000049957 and TRCN0000289044), both part of the TRC library from the Broad Institute. It was only later that we learned that even though the shRNAs had two different clone identifiers, they were in fact the exact same sequence. We hope that this cautionary tale will help others avoid off-target activation of dsRNA sensors such as PKR and improve the rigor and reproducibility of research in this area.

## Materials and Methods

### Cell culture

Breast cancer cell lines (MCF-7 (RRID: CVCL_0031), SK-BR-3 (RRID: CVCL_0033), BT549 (RRID: CVCL_1092), MDA-MB-231 (RRID: CVCL_0062)) and 293T (RRID: CVCL_0063) were obtained from American Type Culture Collection. All cell lines were cultured in Dulbecco’s modified Eagle’s medium (DMEM) (Hyclone) with 10% fetal bovine serum (Bio-Techne), 2 mM glutamine (Hyclone), 0.1 mM nonessential amino acids (Hyclone), 1 mM sodium pyruvate (Hyclone).

### Viral Production and Transduction

Lentivirus was produced by Turbo DNAfection 3000 or LipoFexin (Lamda Biotech) transfection of 293T cells with pCMV-VSV-G, pCMV-ΔR8.2, and pLKO.1-puro (shSCR and shDDX54) or pLKO.1-hygro (shSCR or shADAR) for shRNAs or pLVX-IRES-Hygro for overexpression of DDX54. Virus was harvested 48 hours post-transfection. Cells were transduced with lentivirus for 16 hours in the presence of 10 µg/mL protamine sulfate. The cells were selected with puromycin at 2 µg/mL for two days or 100 mg/mL Hygromycin B for five days.

### Plasmids

The DDX54 overexpression plasmid (pLVX-IRES-Hygro-FH-DDX54) was generated by PCR amplification of DDX54 from pFRT/TO/FLAG/HA-DDX54, a gift from Markus Landthaler (Addgene plasmid # 97060; http://n2t.net/addgene:97060; RRID: Addgene 97060) and ligation into pLVX-IRES-Hygro. The DDX54 wobble mutant was generated by site-directed mutagenesis using PrimeSTAR Max and In-Fusion cloning (Takara). All primers are available in the Supplemental Information. The lentiviral shRNA plasmids for DDX54 were purchased from Millipore-Sigma, TRCN0000049957 and TRCN0000289044. The shSCR plasmid was a gift from the lab of Sheila Stewart, Washington University in St. Louis.

### Immunoblot

Cell pellets were lysed and sonicated in RIPA Buffer (50 mM Tris pH 7.4, 150 mM NaCl, 1% Triton X-100, 0.1% sodium dodecyl sulfate and 0.5% sodium deoxycholate) with 1x HALT Protease Inhibitor (Pierce). Thirty micrograms of protein lysate were resolved on 4-12% TGX Acrylamide Stain-Free gels (Bio-Rad). The Stain-Free gels were imaged prior to transfer to PVDF (Bio-Rad) by TransBlot Turbo (Bio-Rad). The blots were then probed with the appropriate primary antibodies: Primary antibodies: ADAR1 (Santa Cruz, sc-73408; Bethyl Laboratories, A303-883A), DDX54 (Novus Biologicals, NB100-60678), eIF2a (Abcam, ab5369), eIF2a-Ser-51-P (Abcam, ab32157), beta-tubulin (Abcam, catalog no. ab6046, RRID:AB_2210370), cleaved PARP (Cell Signaling Technology, catalog no. 9541, RRID:AB_331426), PKR (Cell Signaling Technology, catalog no. 3072, RRID:AB_2277600), PKR Thr-446-P (Abcam, catalog no. ab32036, RRID:AB_777310). Primary antibodies were detected with horseradish-peroxidase conjugated secondary antibodies (Jackson ImmunoResearch) and detection was carried out with Clarity Western ECL Substrate (Bio-Rad). Densitometry was performed using Image Lab (Bio-Rad). Band intensity was normalized to total protein measured by imaging of the Stain-Free gel.

### Foci Formation Assay

Five thousand cells were plated for each condition in a 10 cm culture dish. After 10 (BT549, MB231 and SK-BR-3) to 20 (MCF-) days the cells were washed briefly with 1x PBS prior to fixation in 100% methanol for 5 min. After drying, the cells were stained with 0.005% Crystal Violet solution containing 25% methanol (Sigma-Aldrich) prior to washing excess stain away with deionized water. The plates were scanned using an ImageScanner III (General Electric). Foci area was calculated using ImageJ.

### PKR Kinase Assay

Recombinant PKR was purified from E. coli as described previously ^25^. The PKR kinase assay was performed using purified PKR and either shDDX54-4 (rGrGrCrCrCrGrGrUrGrUrUrCrArArArGrGrCrArUrCrArUrCrUrCrGrArGrArUrGrArUrGrCrCrUrUr UrGrArArCrArCrCrGrGrGrUrUrUrUrU) or shSCR (rGrGrUrCrCrUrArArGrGrUrUrArArGrUrCrGrCrCrCrUrCrGrCrUrCrGrArGrCrGrArGrGrGrCrGrAr CrUrUrArArCrCrUrUrArGrGrUrUrUrUrU), both synthesized by Integrated DNA Technologies as RNA oligonucleotides. The kinase assay was performed as described previously ^25^.

## Data and Code Availability

Scripts used for data analysis are available here https://github.com/cottrellka/Cottrell-Ryu-et-al-2024. All other data are provided in the main Figures or supplemental Figures and tables.

## Acknowledgements

This work was supported by funding to J.D.W from the National Cancer Institute (R01CA262804), funding to B.L.B from the National Institute of General Medical Sciences (R35GM141262), and funding to K.A.C. from the National Institute on Minority Health and Health Disparities (K99MD016946 and R00MD016946).

