## Supplemental Figures for "Activation of PKR by a short-hairpin RNA"

Department of Biochemistry

Purdue University

201 S University St.

West Lafayette, IN

Jason D. Weber, Ph.D.

Department of Medicine

Division of Molecular Oncology

Washington University School of Medicine

660 South Euclid Avenue

Campus Box 8069

St. Louis, MO 63110 USA

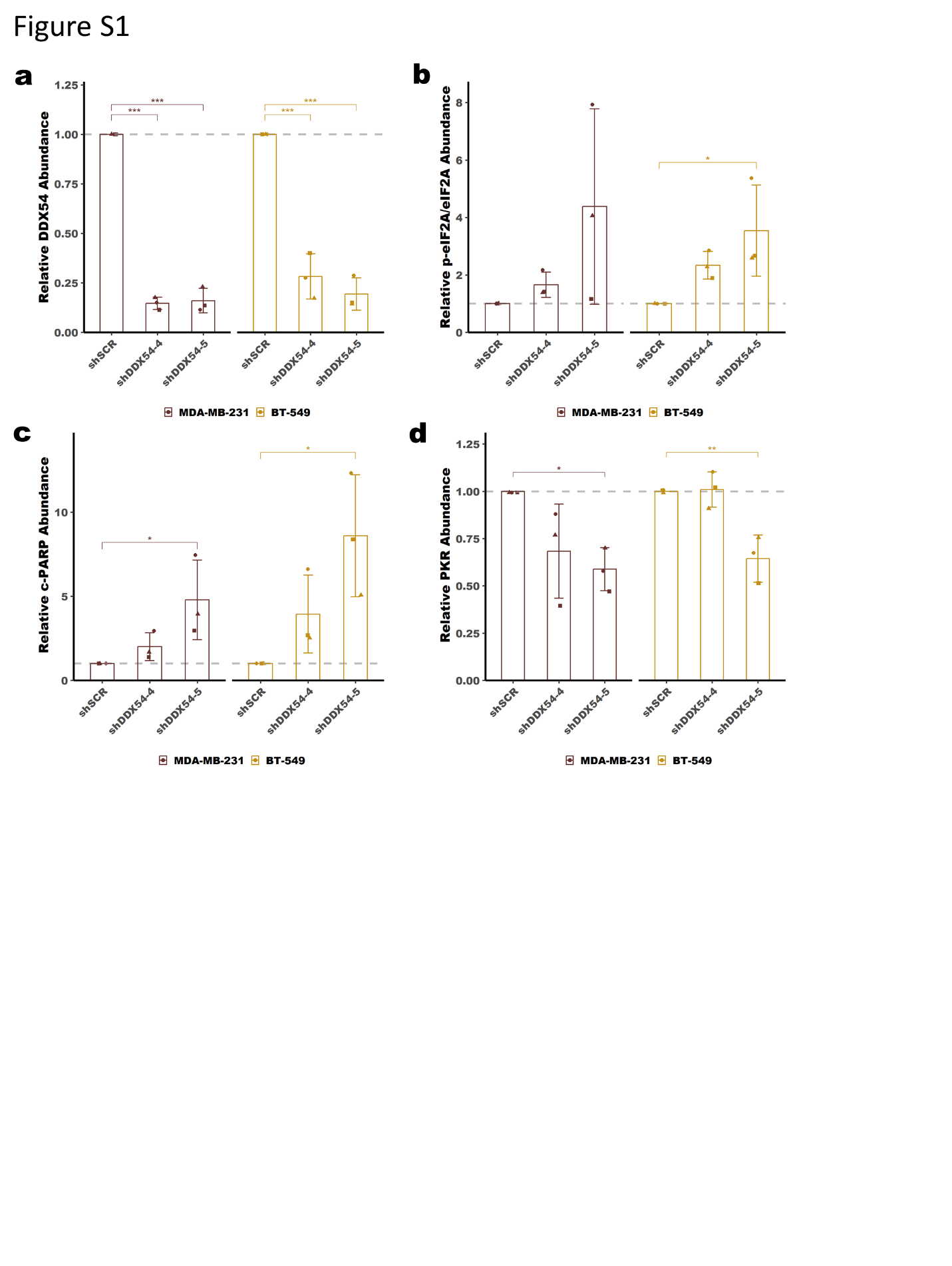

**Supplemental Figure 1:**

**a-d** Quantification of protein abundance from the immunoblot in **Figure 1a**. Bars represent the average of at least three replicates (shown as differently shaped points), error bars are +/- SD. * p <0.05, ** p <0.01, *** p < 0.001. P-values determined by Dunnett’s test.

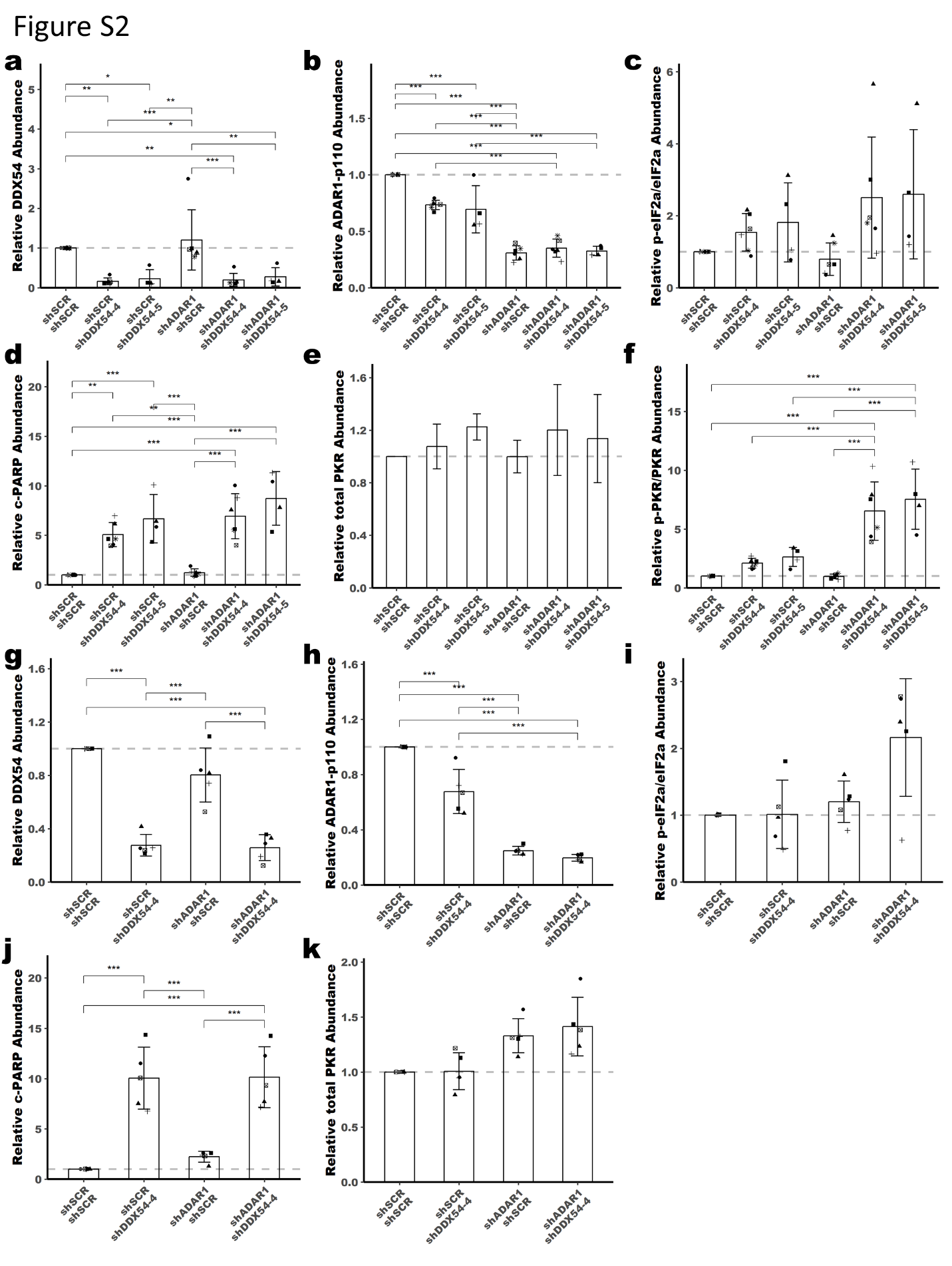

**Supplemental Figure 2:**

**a-f** Quantification of protein abundance from the immunoblot in **Figure 2a**. **g-k** Quantification of protein abundance from the immunoblot in **Figure 2e** Bars represent the average of at least three replicates (shown as differently shaped points), error bars are +/- SD. * p <0.05, ** p <0.01, *** p < 0.001. P-values determined by one-way ANOVA with post-hoc Tukey.

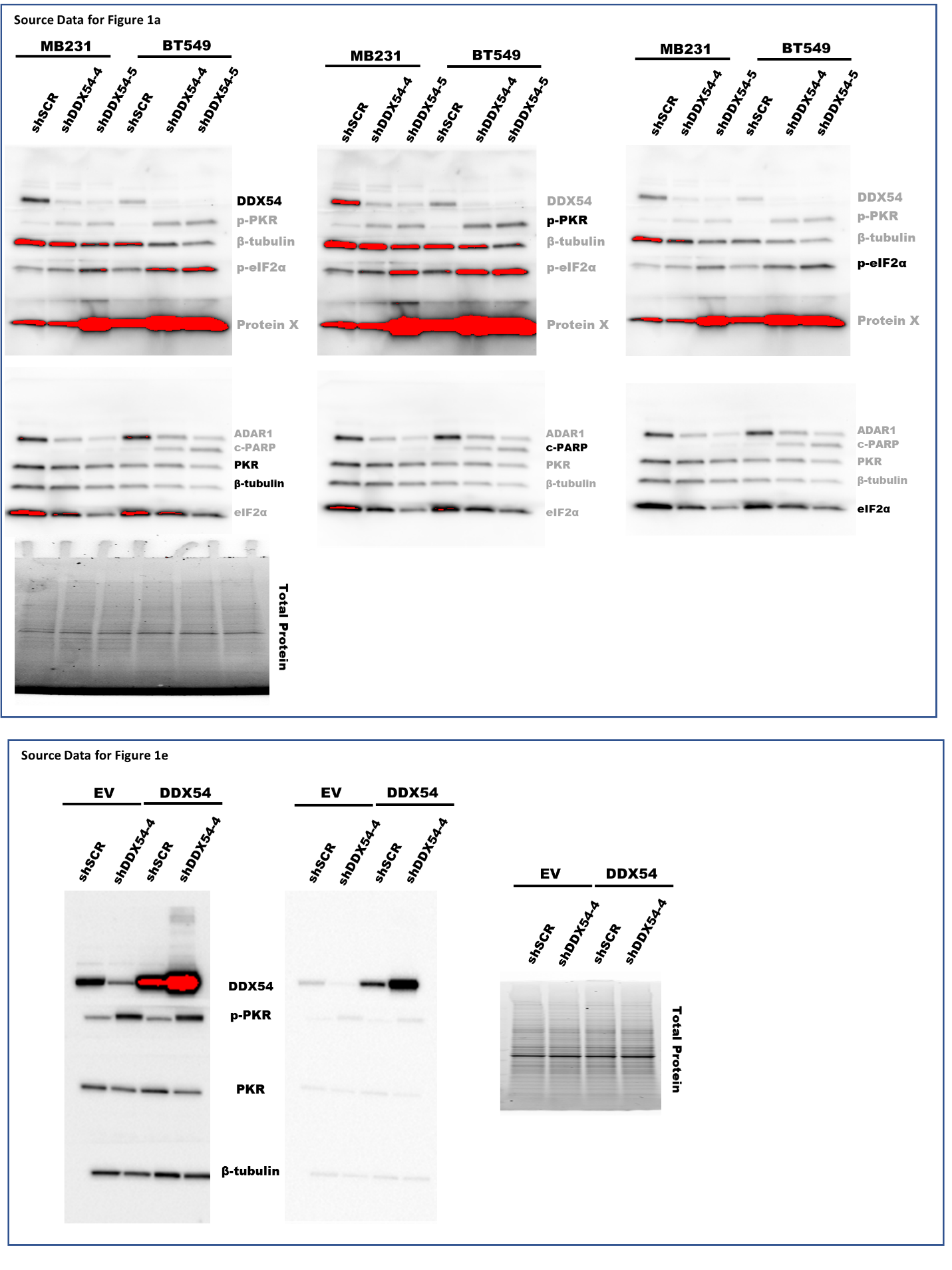

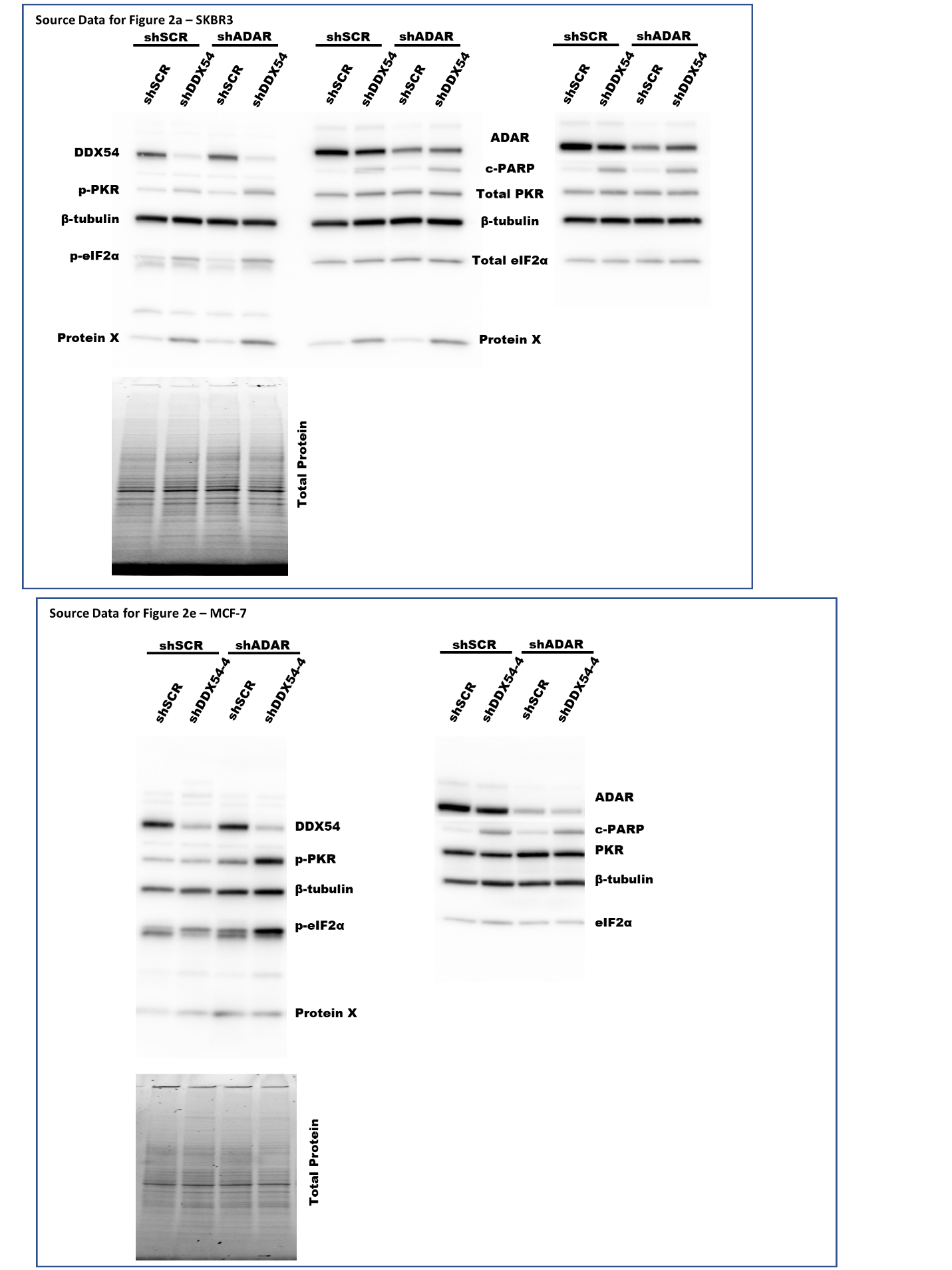
